# How good are pathogenicity predictors in detecting benign variants?

**DOI:** 10.1101/408153

**Authors:** Abhishek Niroula, Mauno Vihinen

**Affiliations:** Protein Structure and Bioinformatics, Department of Experimental Medical Science, Lund University, BMC B13, SE-221 84 Lund, Sweden

**Keywords:** performance assessment, variation interpretation, computational tools, prediction performance, ExAC dataset

## Abstract

Computational tools are widely used for interpreting variants detected in sequencing projects. The choice of computational tools is critical for reliable variant impact interpretation for precision medicine and should be based on systematic performance assessment. The performance of the methods varies widely in different performance assessments, for example due to the sizes of test datasets. To address this issue, we obtained 63,160 common amino acid substitutions (allele frequency ≥1% and <25%) from the Exome Aggregation Consortium (ExAC) database, which contains variants from 60,706 genomes or exomes. We evaluated the specificity, the capability to detect benign variants, for 10 variant interpretation tools. In addition to overall specificity of the tools, we tested their performance for variants in six geographical populations. PON-P2 had the best performance (95.5%) followed by FATHMM (86.4%) and VEST (83.5%). While these tools had excellent performance, the poorest method predicted more than one third of the benign variants to be disease-causing. The results allow choosing reliable methods for benign variant interpretation, for both research and clinical purposes, as well as provide a benchmark for method developers.

**AUTHOR SUMMARY:** In precision/personalized medicine of many conditions it is essential to investigate individual’s genome. Interpretation of the observed variation (mutation) sets is feasible only with computational approaches. We assessed the performance of variant pathogenicity/tolerance prediction programs on benign variants. Variants were obtained from high-quality ExAC database and selected to have minor allele frequency between 1 and 25%. We obtained 63,160 such cases and investigated 10 widely used predictors. Specificities of the methods showed large differences, from 0.64 to 0.96%, thus users of these methods have to be careful when choosing the one(s) they will use. We investigated further the performances on different populations, allele frequencies, separately for males and females, chromosome wise and for population unique and non-unique variants. The ranking of the tools remained the same in all these scenarios, i.e. the best methods were the best irrespective on how the data is filtered and grouped. This is to our knowledge the first evaluation of method performance on benign variants.

## INTRODUCTION

Next Generation Sequencing (NGS) is widely used in clinical diagnosis as well as in population genetics to investigate patterns of genetic variants in healthy individuals. The large numbers of variants, millions per genome in comparison to reference sequences, pose challenges for detecting disease-causing variants. There are on average about 10,000 variants per genome that cause amino acid substitutions [1]. Several databases enable annotation of disease relevance of variants and frequencies among healthy individuals. These include numerous locus specific variation databases (LSDBs) that are curated by experts in the genes and diseases. While LSDBs typically concentrate on individual genes and proteins or diseases, the general databases have much wider scope such as ClinVar [2], Online Mendelian Inheritance in Man (OMIM) [3] and the UniProt Knowledgebase (UniProtKB) [4].

The most harmful variants confer adverse impacts and reduce the fitness of the carrier, and are therefore selected against and removed from the population. On the other hand, the benign variants are tolerated and are inherited through the generations. Therefore, variants occurring at high frequencies in a population are likely benign. Information for variants and their frequencies in various populations are available e.g. in the database of short genetic variations (dbSNP) [5], the 1000 Genomes Project [6], the Exome Sequencing Project (ESP) Exome Variant Server (EVS) [7], and recently in the Exome Aggregation Consortium (ExAC) database [8]. These resources are widely used to filter out likely benign variants as well as for training and testing computational tools. Variants with allele frequencies (AFs) ≥1% are generally assumed to be benign. There are some exceptions e.g. in late onset diseases or due to incomplete penetrance. We are not aware of reliable estimates of such cases. Based on existing literature, the number is so low that it does not affect results based on large scale studies, as in here. Most variants in these databases are rare, for example in the ExAC database, 99% of the variants have AF below 1% [8], and have unknown clinical relevance.

Prediction tools are instrumental for variant effect interpretation in personalized and precision medicine since experimental methods cannot deal with the amounts of variation data generated in sequencing projects. The American College of Medical Genetics and Genomics (ACMG) and the European Society of Human Genetics (ESHG) guidelines recommend using computational predictions as one of several lines of evidence for variant interpretation [9, 10]. Similarly, the joint consensus recommendation for the interpretation of variants in cancer by the Association for Molecular Pathology, American Society of Clinical Oncology, and College of American Pathologists include the use of computational predictions [11].

Numerous computational tools based on different principles have been developed to predict the tolerance and pathogenicity of genetic variants [12-15]. The performance of these tools varies widely [12, 16-19]. Even a minor difference in the performance leads to misinterpretation of large numbers of variants in genome or exome-wide scale. Hence, the choice of the tools is critical for reliable variant interpretation. The assessment of method performance requires benchmark datasets with known outcomes. In this field, such datasets are available at VariBench [20] and VariSNP [21]. Further, the assessment has to be made in a systematic way and reporting the full performance of the analyzed methods [22, 23], which unfortunately is often not the case, especially for commercial products [24]. In addition to pathogenicity/tolerance method assessment, the performance of some other predictor classes have been assessed including alternative splicing [25, 26], protein stability [27, 28], protein solubility [29], and protein localization [30].

A comprehensive predictor assessment requires a benchmark with both positive (showing the effect) and negative (not having an effect) variants. Here, we tested the predictor sensitivity i.e. the capability to recognize variants not having phenotypic effect using the largest available dataset of likely benign variants. Recently, the ExAC database that has been carefully curated and contains quality-controlled data for altogether 60,706 exomes was released [8]. The database contains the overall frequencies of variations across all the individuals as well as the frequencies for several populations. We obtained the common variants from the ExAC database and identified those leading to amino acid substitutions (AASs). In total, 63,160 AASs had AF ≥1% and <25% in at least one of the populations. These AASs are widely considered as benign and therefore were used to assess the performance of the prediction tools. We investigated the performance of 10 widely used prediction methods and found that the best tools are excellent while some others have poor performance.

## MATERIALS AND METHODS

### Variation data

The variation data were obtained from the ExAC database (release 0.3.1) [8] in a Variant Call Format (VCF) file. We identified the variants leading to amino acid substitutions (AASs) by using the annotations from the Variant Effect Predictor (VEP) [31] included in the downloaded VCF file. The amino acid substitutions were further filtered by using the AFs in the whole dataset as well as in different populations. The VCF file contained AFs for various datasets and populations. The adjusted AF (AF for all individuals with genotype quality (GQ) ≥20 and depth (DP) ≥10) as well as the AFs in all geographical populations (African, American, East Asian, Finnish, non-Finnish European, South Asian, and Other) were used in the analysis. In addition, we used the AFs for males and females. Variants having AFs ≥1% and <25% in any of the 9 populations were included to the study. We set an upper threshold of AF to 25%, so that the AFs represented the minor alleles. In total, there are 63,197 variants that meet these criteria. The dataset is available at VariBench (http://structure.bmc.lu.se/VariBench/exac_aas.php).

### Computational predictions

The predictions were obtained from the dbNSFP database (version 3.2a) [32] for several tools. The database contains annotations and predictions for all potential single nucleotide substitution-caused AASs. We obtained the predictions for Combined Annotation Dependent Depletion (CADD) [33], Functional Analysis through Hidden Markov Models (FATHMM) [34], Likelihood Ratio Test (LRT) [35], MutationAssessor [36], MetaLR [37], MetaSVM [37], MutationTaster2 [38], PolyPhen-2 [39], Protein Variation Effect Analyzer (PROVEAN) [40], Sorting Intolerant From Tolerant (SIFT) [41], and Variant Effect Scoring Tool (VEST) [42]. If there were multiple predictions for a variant from the same tool, we took the most frequent classification. If two classes were equally frequent, then the classification was considered as ambiguous. In addition, we obtained predictions for PON-P2 [18] by using the tool’s Application Programming Interface (API).

### Training datasets

Training datasets were obtained for FATHMM, MetaLR, MetaSVM, PolyPhen-2, VEST, and PON-P2 and cases in them were excluded from assessment of those tools. Since no variations were left for Meta-LR and Meta-SVM after excluding the training data, we could not evaluate these methods.

### Common variants

Variants with AF ≥1% and <25% in a specific population are considered as common for that population. This criterion was used to obtain 10 subsets of variation data. For the six geographical populations: African/African American (AFR), Latino (AMR), East Asian (EAS), Finnish (FIN), Non-Finnish European (NFE), and South Asian (SAS), the datasets were further partitioned into population-specific unique and non-unique datasets. The unique dataset contains variants with AF ≥1% and <25% in the specific population but <1% in all other populations and the non-unique dataset consists of the remaining variants. For example, the variants with AF ≥1% and <25% in AFR population are indicated as common variants for AFR population. From those, the variants with AF <1% in all the five other geographical populations are unique variants for the AFR population. The remaining common variants in the AFR population are non-unique variants.

To exclude misclassified pathogenic variants in the dataset filtered with the AF threshold, we obtained from ClinVar all the 24,232 variants that lead to AASs and were annotated as pathogenic or likely pathogenic (13 July 2018) [2]. There were 37 variants which had AF ≥1% and <25%, some of which had been used for predictor training: FATHMM (14 variants), PON-P2 (14), PolyPhen-2 (4), and VEST (6). The reason at least for some of these variants to be included into the training datasets is that more data may have accumulated to reclassify variants after the methods were trained.

### Performance comparison

Except for CADD and VEST, the investigated methods classify the variants into harmful and benign. We used these classifications for the method performance assessment. For CADD, we classified the variants based on the phred-like score with a cutoff 20, below which the variants were classified as benign and otherwise harmful, as suggested by the authors. For VEST, we classified the variants based on the VEST score with a cut-off 0.5, below which the variants were classified as benign and otherwise harmful. The terms *deleterious, damaging, probably damaging, possibly damaging, disease-causing, functional*, and *pathogenic* were all considered to be harmful and the terms *tolerated, benign, neutral, non-functional*, and *polymorphism* were all considered to be benign. MutationTaster2 provides automatic annotations for harmful and benign variants based on annotations in variation databases and predicts the impacts for others. In this study, the automatic annotations of MutationTaster2 were excluded to test the actual prediction capability of the tool. PON-P2 and LRT classify variants into three classes, the third class being variants of unknown significance. The variants classified as unknown were excluded.

Several measures are needed to describe the overall performance of prediction methods [22, 23]. Since we investigated only one type of variants, the benign ones, it was possible to calculate only a single measure, the specificity. Specificity is the proportion of correctly predicted benign variants,

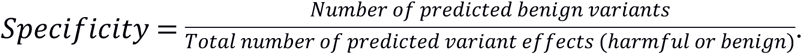

The scores can be multiplied by 100 to show results in percentages.

## RESULTS

### Specificity of tolerance predictors

To assess the quality of variant pathogenicity/tolerance prediction methods we collected from the ExAC database all variants that had AF ≥1% and <25%. Because of their high frequency, these variants are usually considered to be neutral and were used in here to assess the specificity of prediction methods. The predictions for 9 tools were collected from the dbNSFP database [32]. For PON-P2 [18], we run the predictions using the Application Programming Interface. The tools are based on different principles and include those based on evolutionary information only, LRT [35], PROVEAN [40], and SIFT [41], and those combining different types of features, CADD [33], FATHMM [34], MutationAssessor [36], MutationTaster2 [43], PolyPhen2 [39], PON-P2 [44], and VEST [42]. Most of the investigated tools have been trained with known disease-causing and benign variants. The methods that use only sequence conservation information have not been trained. If variants used for training are used for assessing the methods, the obtained performance measures are likely inflated [16, 22, 45]. Hence, we excluded the training datasets for FATHMM, PON-P2, PolyPhen-2, and VEST. The remaining tools were either not trained or the training datasets were not available.

The performances of some of these tools have been assessed previously several times, however not with this kind of high-quality and large dataset for benign variants. It is important both in research and clinical practice to be able to sort out variants that have no relevance for the condition under investigation. The specificities of the methods range from 0.63 for SIFT and 0.64 for MutationTaster2 to 0.96 for PON-P2 (Table 1). FATHMM and VEST have the second and third highest performance i.e. 0.86 and 0.84, respectively. It should be noted that variants are classified into three classes by PON-P2 and two classes by FATHMM, and VEST predicts continuous probabilities. For VEST, we classified the variants into two classes using a cutoff of 0.5. PON-P2 had the highest proportion of unclassified variants, however with far better specificity compared to the other tools (Fig. 1 and Table 1). To assess the performance of tools for variants with different AFs, we divided the dataset into groups based on adjusted AF on the whole dataset. The predictor performance is higher for variants with higher AFs for all the tools (Fig. 1). The differences between the AF bins are the smallest for PON-P2 and FATHMM while several other methods had very strong correlation between specificity score and allele frequency.

**Table 1.**
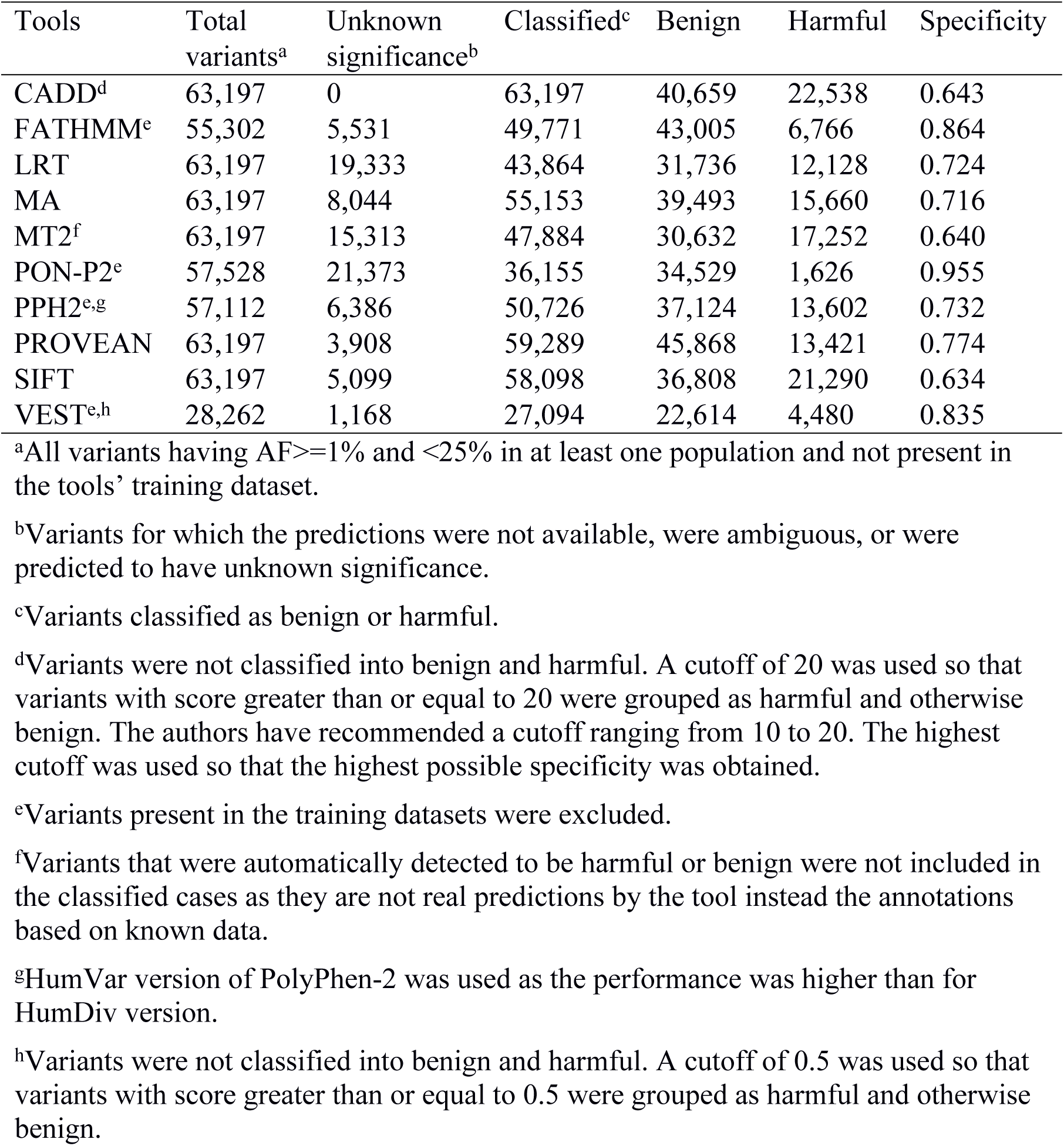
Specificities of variant interpretation tools on all variants in the dataset (AF ≥1% and <25%).

**Figure 1.**
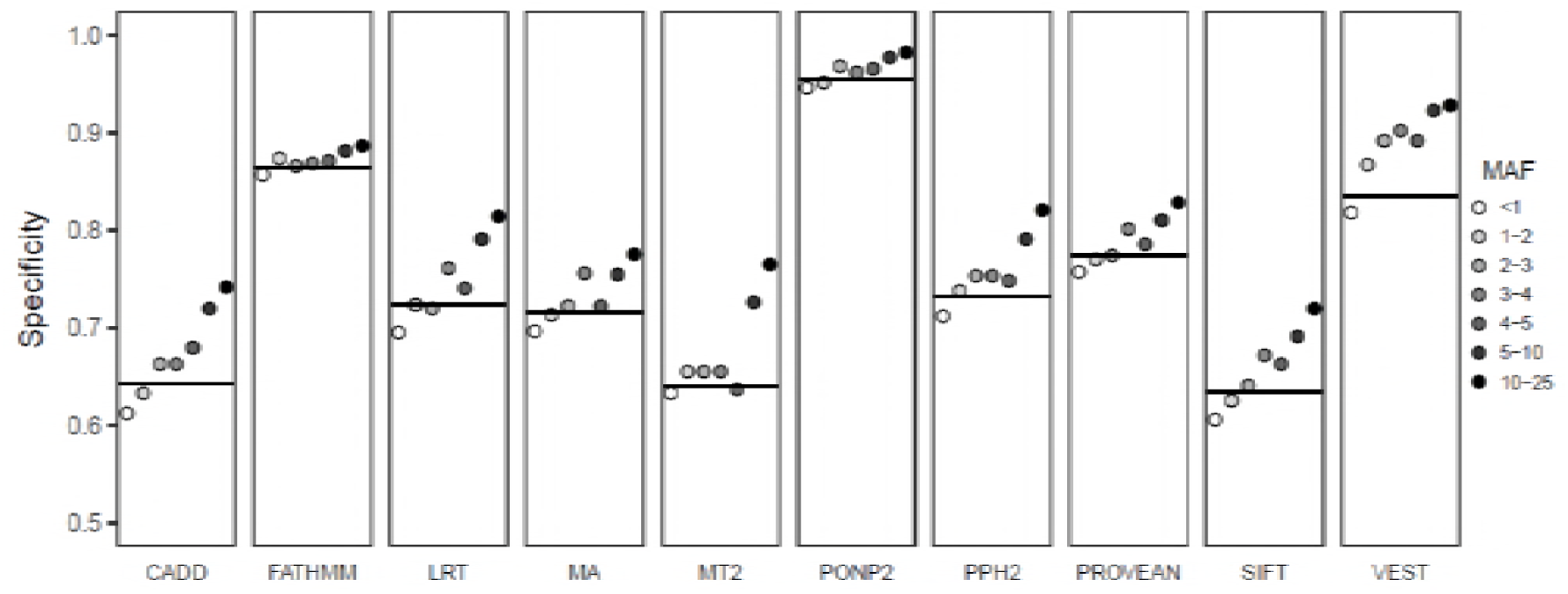
Performance of variant tolerance predictors. Specificities of 10 prediction tools for variants with different AFs. The black horizontal line indicates performance for all variants (AF ≥1% and <25%). The variants with AF <1% have low AF in the whole dataset but have higher AF in at least one of the populations. MA, MutationAssessor; MT2, MutationTaster2; PPH2, PolyPhen-2.

### Population-specific performance

ExAC database contains information for the ethnic origin of the individuals. Thus, in addition to the general performance, we were able to analyze also ethnicity-based assessment. The same three tools, i.e. PON-P2, FATHMM, and VEST showed the highest specificities also on the data for the populations (Fig. 2). The methods, however, show somewhat different performances for different populations. PON-P2 and FATHMM have small performance difference between the populations while VEST has bigger performance differences. Interestingly, all the tools have the lowest specificity for the Finnish population. This is presumably because the small, and in the past closed population has passed through a narrow bottleneck some 300 years ago during which certain unique alleles were highly enriched.

**Figure 2.**
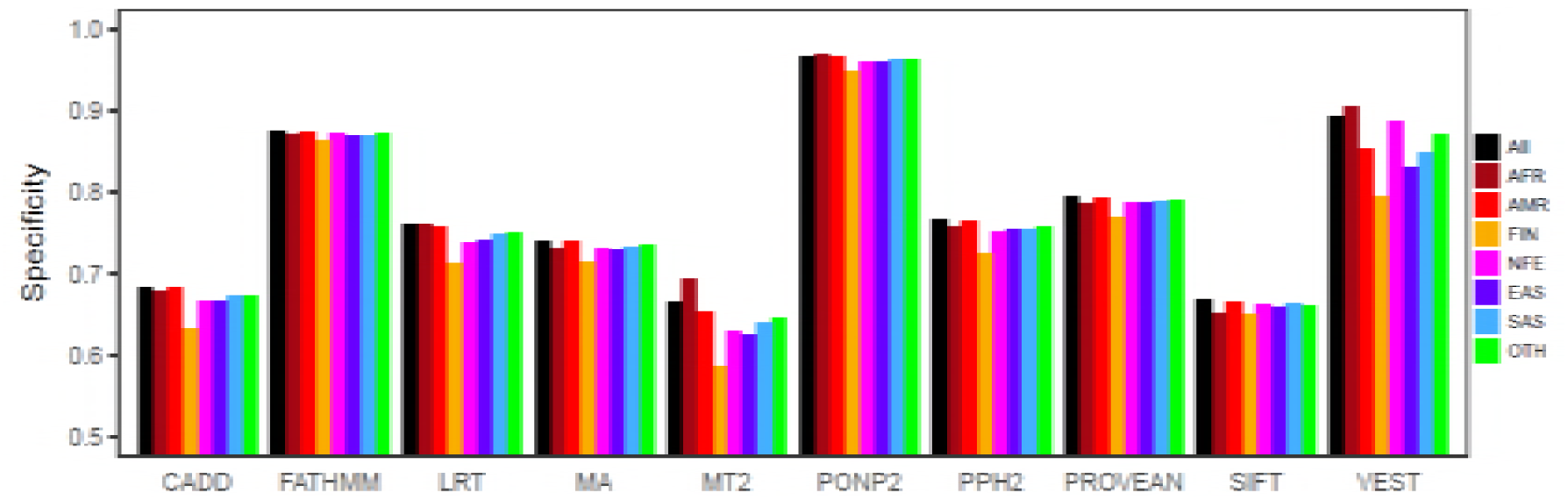
Performance of variant tolerance predictors for variants in ethnic groups. Specificities of prediction tools for common variants (AF ≥1% and <25%) in different populations. AFR, African; AMR, American; EAS, East Asian; FIN, Finnish; NFE, Non-Finnish European; OTH, Other; SAS, South Asian; MA, MutationAssessor; MT2, MutationTaster2; PPH2, PolyPhen-2.

Next, we identified population-specific common variants which have AF ≥1% and <25% in one population but have AF <1% in all the other populations. These are referred to as population-specific unique variants and the remaining variants for the population are referred to as non-unique variants. The proportions of unique variants vary in the populations, ranging from 6.8% in European population (excluding Finnish) to 62.4% in the African population (Table 2). The tools showed lower specificities for the unique variants than for the non-unique variants in the populations (Table 3). The lowest performance is seen for the unique variants in the Finnish population. Performance differences vary largely depending on the tools and the populations. The performance differences between the unique and non-unique variants are the lowest in the African population and the highest in the Finnish population. With respect to the tools, the differences are the lowest for FATHMM (ranging from 1.3 to 4.1%) and PON-P2 (ranging from 1.2 to 8.0%).

**Table 2.** Proportion of unique variants in the populations.

| Population | Total variants<sup>a</sup> | Unique variants<sup>b</sup> | Proportion of<br>unique variants |
| --- | --- | --- | --- |
| AFR | 35,553 | 22,197 | 0.624 |
| AMR | 17,651 | 1,829 | 0.104 |
| EAS | 14,750 | 4,845 | 0.328 |
| FIN | 18,804 | 3,306 | 0.176 |
| NFE | 18,335 | 1,240 | 0.068 |
| OTH | 19,471 | 640 | 0.033 |
| SAS | 18,167 | 3,396 | 0.187 |

<sup>a</sup>All variants having AF $\geq 1\%$ and $< 25\%$ in the population.
<sup>b</sup>Variants having AF $\geq 1\%$ and $< 25\%$ in the specific population but $< 1\%$ in all other populations.

**Table 3.**
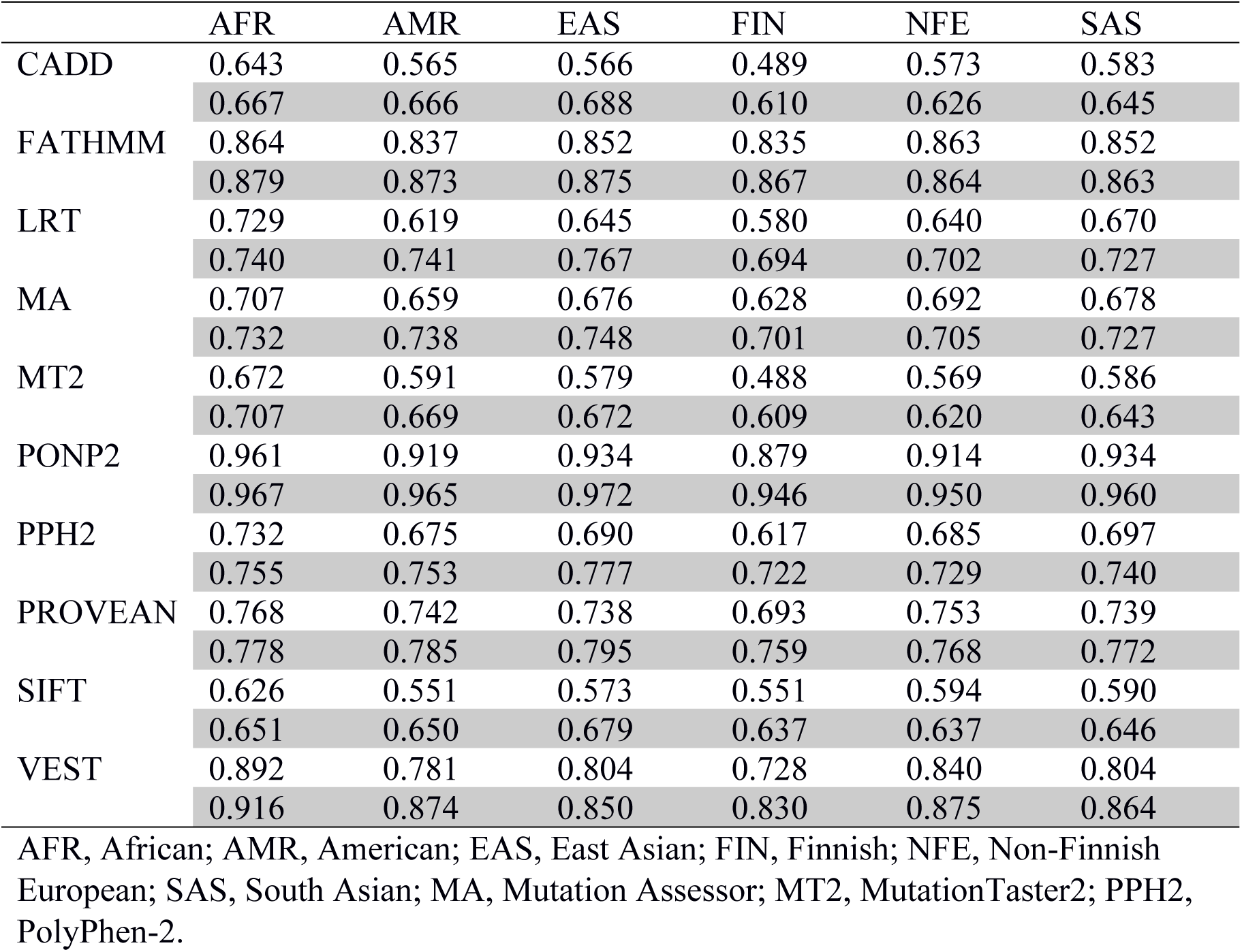
Specificities of tools for unique and non-unique variants in six populations. The performance scores with grey background indicate specificities for non-unique variants in the populations.

|  | AFR | AMR | EAS | FIN | NFE | SAS |
| --- | --- | --- | --- | --- | --- | --- |
| CADD | 0.643 | 0.565 | 0.566 | 0.489 | 0.573 | 0.583 |
|  | 0.667 | 0.666 | 0.688 | 0.610 | 0.626 | 0.645 |
| FATHMM | 0.864 | 0.837 | 0.852 | 0.835 | 0.863 | 0.852 |
|  | 0.879 | 0.873 | 0.875 | 0.867 | 0.864 | 0.863 |
| LRT | 0.729 | 0.619 | 0.645 | 0.580 | 0.640 | 0.670 |
|  | 0.740 | 0.741 | 0.767 | 0.694 | 0.702 | 0.727 |
| MA | 0.707 | 0.659 | 0.676 | 0.628 | 0.692 | 0.678 |
|  | 0.732 | 0.738 | 0.748 | 0.701 | 0.705 | 0.727 |
| MT2 | 0.672 | 0.591 | 0.579 | 0.488 | 0.569 | 0.586 |
|  | 0.707 | 0.669 | 0.672 | 0.609 | 0.620 | 0.643 |
| PONP2 | 0.961 | 0.919 | 0.934 | 0.879 | 0.914 | 0.934 |
|  | 0.967 | 0.965 | 0.972 | 0.946 | 0.950 | 0.960 |
| PPH2 | 0.732 | 0.675 | 0.690 | 0.617 | 0.685 | 0.697 |
|  | 0.755 | 0.753 | 0.777 | 0.722 | 0.729 | 0.740 |
| PROVEAN | 0.768 | 0.742 | 0.738 | 0.693 | 0.753 | 0.739 |
|  | 0.778 | 0.785 | 0.795 | 0.759 | 0.768 | 0.772 |
| SIFT | 0.626 | 0.551 | 0.573 | 0.551 | 0.594 | 0.590 |
|  | 0.651 | 0.650 | 0.679 | 0.637 | 0.637 | 0.646 |
| VEST | 0.892 | 0.781 | 0.804 | 0.728 | 0.840 | 0.804 |
|  | 0.916 | 0.874 | 0.850 | 0.830 | 0.875 | 0.864 |

AFR, African; AMR, American; EAS, East Asian; FIN, Finnish; NFE, Non-Finnish European; SAS, South Asian; MA, Mutation Assessor; MT2, MutationTaster2; PPH2, PolyPhen-2.

As the tools have lower performances for unique variants, we investigated the frequencies of unique variants and those that were not unique (i.e. non-unique). Most unique variants have low AF, between 1% and 5%, while the proportions of non-unique variants with different AFs are similar (Fig. 3). Since many predictors have been trained with variants with high allele frequencies, the lower specificities for unique variants could be due to disparity in the frequencies. To control bias due to frequency, we compared the performance of the tools for unique and non-unique variants with AF in the same range (i.e. 1-5%) in each population. The comparison showed that the tools indeed have poorer performance for unique variants than for non-unique variants (Table 4).

**Table 4.** Specificities of tools for unique and non-unique variants with AF ≥1% and <5% in the populations. The scores with grey background indicate specificities for the non-unique variants.

|  | AFR | AMR | EAS | FIN | NFE | SAS |
| --- | --- | --- | --- | --- | --- | --- |
| PON-P2 | 0.961 | 0.918 | 0.934 | 0.879 | 0.914 | 0.934 |
|  | 0.967 | 0.965 | 0.972 | 0.946 | 0.95 | 0.96 |
| SIFT | 0.626 | 0.551 | 0.572 | 0.551 | 0.595 | 0.59 |
|  | 0.651 | 0.65 | 0.679 | 0.637 | 0.637 | 0.646 |
| PPH2 | 0.732 | 0.674 | 0.69 | 0.616 | 0.685 | 0.697 |
|  | 0.755 | 0.753 | 0.778 | 0.722 | 0.729 | 0.74 |
| LRT | 0.729 | 0.618 | 0.645 | 0.579 | 0.639 | 0.67 |
|  | 0.74 | 0.741 | 0.767 | 0.694 | 0.702 | 0.727 |
| MT2 | 0.672 | 0.591 | 0.579 | 0.487 | 0.569 | 0.586 |
|  | 0.707 | 0.669 | 0.672 | 0.609 | 0.62 | 0.643 |
| MA | 0.707 | 0.659 | 0.675 | 0.627 | 0.692 | 0.678 |
|  | 0.732 | 0.738 | 0.748 | 0.701 | 0.705 | 0.727 |
| FATHMM | 0.864 | 0.836 | 0.853 | 0.835 | 0.863 | 0.852 |
|  | 0.879 | 0.873 | 0.875 | 0.867 | 0.864 | 0.863 |
| PROVEAN | 0.768 | 0.742 | 0.737 | 0.692 | 0.753 | 0.739 |
|  | 0.778 | 0.785 | 0.795 | 0.759 | 0.768 | 0.772 |
| CADD | 0.643 | 0.564 | 0.566 | 0.489 | 0.573 | 0.583 |
|  | 0.667 | 0.666 | 0.688 | 0.61 | 0.626 | 0.645 |
| VEST | 0.892 | 0.78 | 0.803 | 0.728 | 0.838 | 0.804 |
|  | 0.915 | 0.874 | 0.85 | 0.829 | 0.874 | 0.863 |

AFR, African; AMR, American; EAS, East Asian; FIN, Finnish; NFE, Non-Finnish European; SAS, South Asian; MA, Mutation Assessor; MT2, MutationTaster2; PPH2, PolyPhen-2.

**Figure 3.**
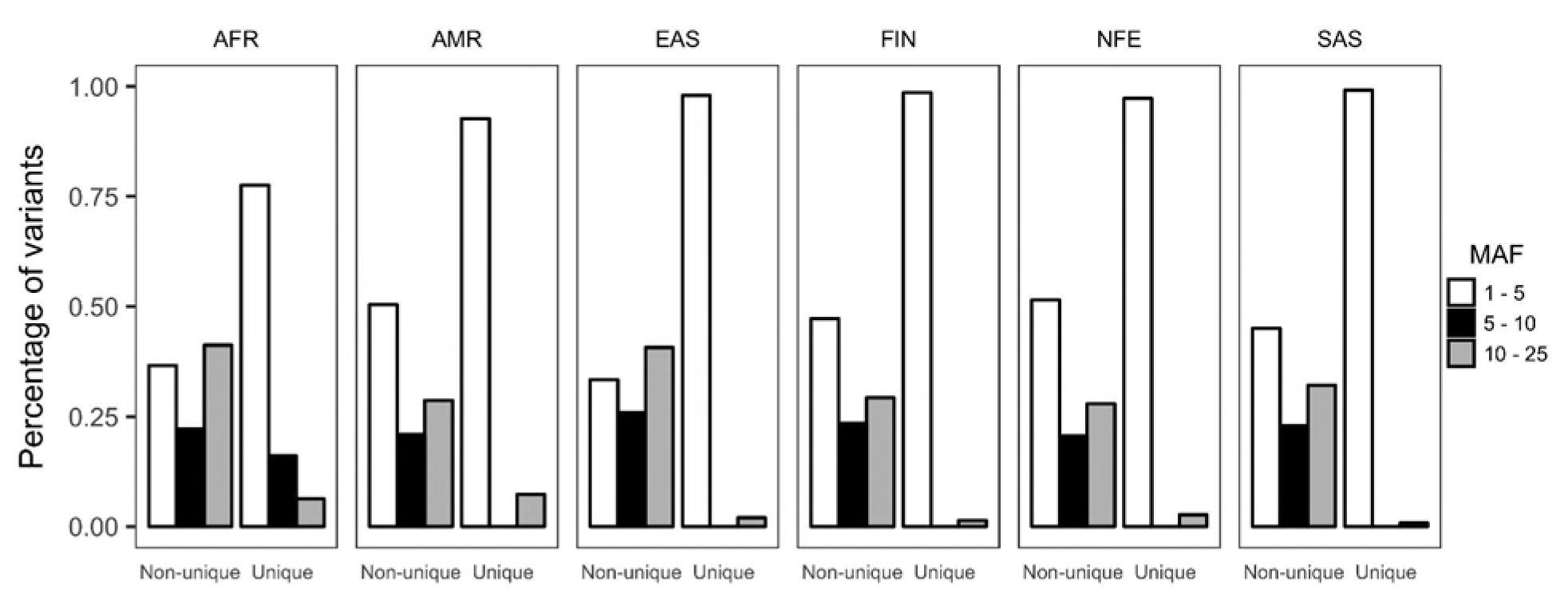
Proportion of unique and non-unique variants with different minor allele frequencies in different populations. AFR, African; AMR, American; EAS, East Asian; FIN, Finnish; NFE, Non-Finnish European; SAS, South Asian.

### Effects of the sex and chromosomal location on prediction performance

Finally, we evaluated the performance for variants from males and females in the populations. No differences were observed in predictor performance. Most of the variants in these two datasets are overlapping. The proportions of unique variants in male (AF ≥1% in male but <1% in female) and female (AF ≥1% in female but <1% in male) populations are 5.6% and 16.9%, respectively (Table 5). The performance for unique variants in male is lower than for the common variants and unique variants in female (Fig. 4).

**Table 5.** Chromosome-wide numbers of variants with AF ≥1% and <25% in male and female populations.

| Chromosome | Variants in female<sup>a</sup> | Variants in male | Ratio of variants in male to female |
| --- | --- | --- | --- |
| 1 | 2,301 (391) | 2,008 (103) | 0.873 (0.263) |
| 2 | 1,420 (234) | 1,268 (77) | 0.893 (0.329) |
| 3 | 1,100 (172) | 981 (54) | 0.892 (0.314) |
| 4 | 847 (147) | 754 (53) | 0.890 (0.361) |
| 5 | 882 (138) | 774 (34) | 0.878 (0.246) |
| 6 | 1,514 (174) | 1,414 (67) | 0.934 (0.385) |
| 7 | 1,039 (204) | 903 (61) | 0.869 (0.299) |
| 8 | 704 (152) | 574 (26) | 0.815 (0.171) |
| 9 | 850 (133) | 760 (40) | 0.894 (0.301) |
| 10 | 881 (126) | 785 (33) | 0.891 (0.262) |
| 11 | 1,409 (238) | 1,239 (70) | 0.879 (0.294) |
| 12 | 1,024 (157) | 907 (47) | 0.886 (0.299) |
| 13 | 316 (37) | 289 (11) | 0.915 (0.297) |
| 14 | 636 (108) | 561 (36) | 0.882 (0.333) |
| 15 | 731 (113) | 661 (44) | 0.904 (0.389) |
| 16 | 1,061 (192) | 936 (62) | 0.882 (0.323) |
| 17 | 1,191 (228) | 1,030 (59) | 0.865 (0.259) |
| 18 | 341 (56) | 302 (18) | 0.886 (0.321) |
| 19 | 1,951 (372) | 1,670 (100) | 0.856 (0.269) |
| 20 | 581 (105) | 500 (29) | 0.861 (0.276) |
| 21 | 277 (45) | 245 (12) | 0.884 (0.267) |
| 22 | 557 (92) | 488 (19) | 0.876 (0.207) |
| X | 428 (110) | 354 (31) | 0.827 (0.282) |
| Y | NA | 3 (3) | NA |
| Total | 22,041 (3724) | 19,406 (1089) | 0.880 (0.292) |
| Percentage of unique variants | 16.9 | 5.6 |  |

<sup>a</sup>The values in brackets are for the unique variants.

**Figure 4.**
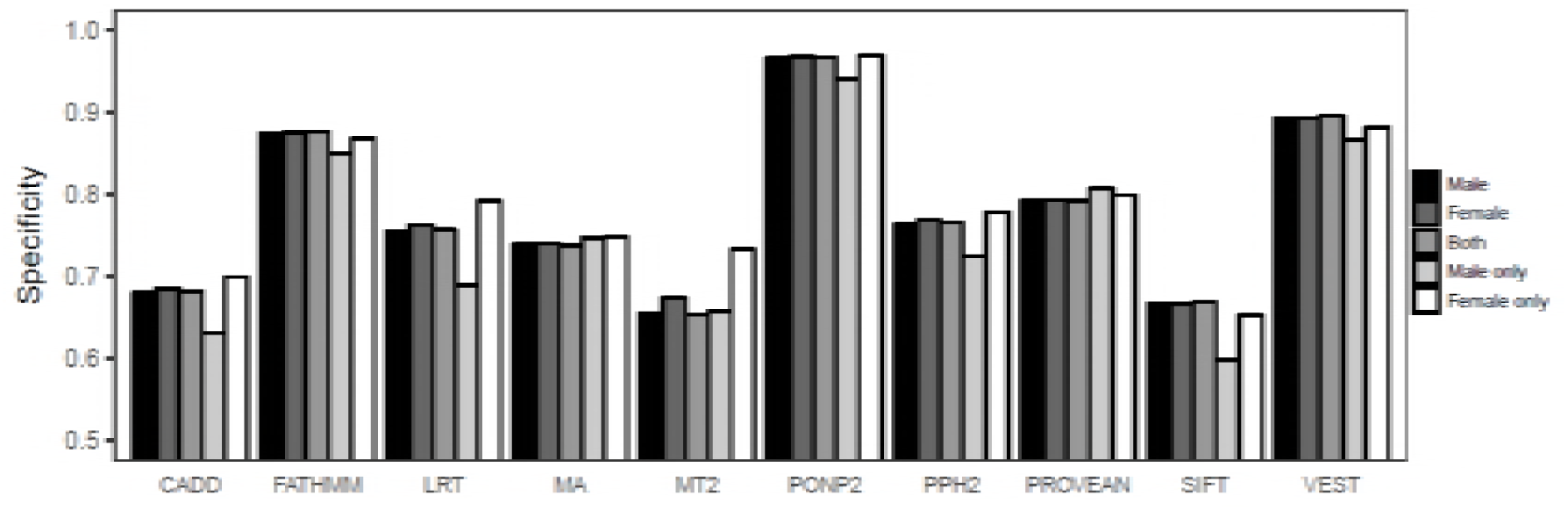
Performance of variant tolerance predictors for variants in males and females.

To assess the influence of variants in sex chromosomes for the lower performance of tools for unique variants in males, we examined the proportions of variants for females and males in all chromosomes. As there were only 3 variants in Y chromosome we could not investigate performance for variants in this chromosome. In the remaining chromosomes, the ratio of unique variants in males to females range from 0.17 to 0.39, with a median of 0.30. The ratio is 0.28 in the X chromosome, i.e. very close to the median (Table 5). The tools show only minor differences in the specificities for variants in different chromosomes (Fig. 5).

**Figure 5.**
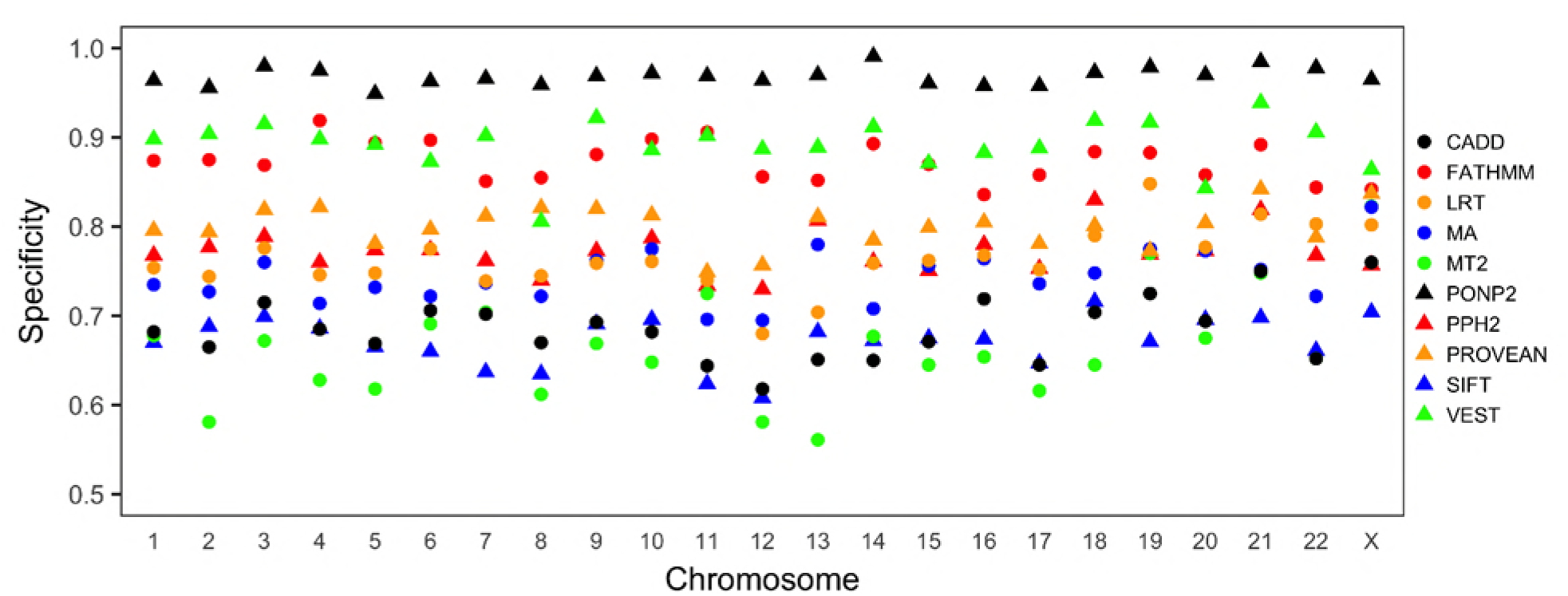
Chromosome-wise performance of tools. Variants in chromosome Y were excluded because there were only 3 variants. MA, Mutation Assessor; MT2, MutationTaster2; PPH2, PolyPhen-2.

## DISCUSSION

Performance comparison of the computational tools enables choosing the most reliable methods. Critical Assessment of Genome Interpretation (CAGI, https://genomeinterpretation.org/) is a community wide effort to assess variant interpretation tools and approaches in the form of competitions [46]. In addition, performance of the tools has been tested by the developers as well as independent researchers. Since some predictors are frequently updated and new ones are developed, they should be assessed regularly [13]. Large datasets of both positive and negative classes are required to assess the performance comprehensively. Due to the lack of a large dataset of disease-causing variations that does not overlap with the training datasets used by the method developers, we could not assess the true positive and false negative rates for the tools. Although several performance measures are required to describe the overall performances of prediction methods [22, 23], we could only compare specificities of the tools, i.e. the capabilities of the tools to detect benign variants. We used the common variants from the ExAC database and the variants predicted to be neutral were considered as correct predictions and those predicted to be disease-related as false negatives. The large size of the ExAC database lends strength for the analysis.

Many tools have been trained with disease-causing and likely benign variants. In most cases, the benign variants have been selected based on their allele frequencies in general population(s). The common variants are considered as benign and the tools have been benchmarked against them. In some rare cases disease-related variants can have high frequency at least in some populations (e.g. sickle cell anemia HbS allele). However, such cases are very rare and are not considered to affect statistics when using large number of cases, as in here.

The analysis of burden of the harmful variants revealed that most harmful variants have extremely low AFs [47]. However, benign variants can have equally low AFs as harmful ones. Performance assessments of tools with variants with all AFs for both harmful and benign variants are desirable; however, such dataset does not exist. In this study, we defined variants with AF ≥1% and <25% as benign variants. The upper limit of 25% was set so that the variant allele analyzed is a minor allele. Although performance evaluation of prediction tools on such common variants may overestimate specificities of the tools, validated benign variants with low AF values are rare. Our results show that specificities increase with AF and have similar trend for all the tools (Fig. 1). Therefore, assessments using the common variants provide useful comparison of the performance of predictors.

Our results show that the performances of tools in detecting the benign variants vary widely. The specificities of the tools ranged from 63.4% to 95.5% (Table 1). PON-P2 [18] had the best performance while MutationTaster2 [38], SIFT [41], and CADD [33] showed the poorest specificities. MutationTaster2 directly annotates the variants as disease-causing or benign based on the dbSNP [5], The 1000 Genomes Project [6], ClinVar [48], and HGMD [49] data. We excluded such automatic annotations in this study to compare the predictive performance of MutationTaster2.

In addition to the specificities of the tools, we also compared the performance on variants common in different geographical populations. All the methods showed performance differences for populations, the lowest specificity was achieved for the variants in the Finnish population (Fig. 2). The variants that were unique in specific populations (AF ≥ 1% and < 25% in the specific population but AF < 1% in all other populations) were more difficult to predict. The tools showed from slightly to markedly lower performance for these variants (Tables 3 and 4). Most of the unique variants had AFs < 5% (Fig. 3). To investigate the possibility of the performance associated with low AF, we compared the performance for the unique variants and the non-unique variants (those with AF ≥ 1% in more than one population) with AF < 5% in the same population. The comparison showed that the specificities were slightly poorer for the unique variants than for the non-unique variants. Differences in the performance on chromosome-wide analysis were very small for all the tools (Fig. 5).

The methods showed very broad spectrum of performances; thus, it is important for the end-users in research as well as in precision medicine to pick a reliable one. Our results enable comparison of the tools and choosing the most reliable ones for variation interpretation.

## ACKNOWLEDGEMENTS

We thank Rachel Karchin and Christopher Douville for providing the training dataset for VEST.

